# Transmembrane 163 (TMEM163) protein effluxes zinc

**DOI:** 10.1101/692525

**Authors:** Vanessa B. Sanchez, Saima Ali, Adrian Escobar, Math P. Cuajungco

**Affiliations:** Department of Biological Science, California State University Fullerton, CA, USA 92831; Center for Applied Biotechnology Studies, California State University Fullerton, CA, USA 92831

**Author notes:** **To whom correspondence should be addressed**: Dr. Math P. Cuajungco, Department of Biological Science, California State University Fullerton, 800 North State College Blvd, Fullerton, CA 92831.

**Keywords:** SV31, zinc transporter, cation diffusion facilitator, single nucleotide polymorphism, ZnT protein

## Abstract

Recent investigations of rodent Tmem163 suggest that it binds to and transports zinc as a dimer, and that alanine mutagenesis of its two species-conserved aspartate (D123A/D127A) residues proposed to bind zinc perturbs protein function. Direct corroboration, however, is lacking whether it is an influx or efflux transporter in cells. We hypothesized that human TMEM163 is a zinc effluxer based on its predicted protein characteristics. We used cultured human cell lines that either stably or transiently expressed TMEM163, and pre-loaded the cells with zinc to determine transport activity. We found that TMEM163-expressing cells exhibited significant reduction of intracellular zinc levels as evidenced by two zinc-specific fluorescent dyes and radionuclide zinc-65. The specificity of the fluorescence signal was confirmed upon treatment with TPEN, a high-affinity zinc chelator. Multiple sequence alignment and phylogenetic analyses showed that TMEM163 is related to distinct members of the cation diffusion facilitator (CDF) protein family. To further characterize the efflux function of TMEM163, we substituted alanine in two homologous aspartate residues (D124A/D128A) and performed site-directed mutagenesis of several conserved amino acid residues identified as non-synonymous single nucleotide polymorphism (S61R, S95C, S193P, and E286K). We found a significant reduction of zinc efflux upon cellular expression of D124A/D128A or E286K protein variant when compared with wild-type, suggesting that these particular amino acids are important for normal protein function. Taken together, our findings demonstrate that TMEM163 effluxes zinc and should now be designated ZNT11, as a new member of the mammalian CDF family of zinc efflux transporters.

---

Abnormal extracellular zinc accumulation leading to cytosolic zinc overload has been shown to result in cytotoxicity *in vitro* and *in vivo*, which can be rescued by zinc-specific chelators (Choi *et al.*, 1988; Koh and Choi, 1994; Cuajungco and Lees, 1996; Koh *et al.*, 1996; Cuajungco and Lees, 1998; Kukic *et al.*, 2014). Many tissues and organs, therefore, have redundant defense mechanisms to strictly maintain zinc homeostasis in order to prevent the cytotoxic effect of intracellular zinc elevation. Zinc homeostasis in mammalian cells is achieved by a tissue-specific and highly conserved zinc transporter protein families called ZNT (SLC30) and Zrt- and Irt-like protein (ZIP; SLC39) (Eide, 2006; Colvin *et al.*, 2010; Kambe *et al.*, 2014). The ZNT proteins are known to efflux zinc from the cytoplasm into the extracellular milieu and into the lumen of membrane-bound or vesicular compartments. The ZIP proteins, meanwhile, influx zinc from the outside of the cell into the cytoplasm and from the lumen of membrane-bound or vesicular compartments into the cytoplasm (Eide, 2006; Colvin *et al.*, 2010; Kambe *et al.*, 2014). In addition to ZNTs, the cells have secondary defense mechanisms to prevent cytoplasmic zinc overload by buffering zinc through metalloproteins, metallothionein (MT), glutathione, and amino acids like histidine and cysteine (Itoh *et al.*, 1983; Aiken *et al.*, 1992; Horn *et al.*, 1995).

Zinc dyshomeostasis has been implicated in several human diseases such as schizophrenia (Scarr *et al.*, 2016), age-related macular degeneration (Smailhodzic *et al.*, 2014), Alzheimer’s disease (Cuajungco and Lees, 1997a; b; Danscher *et al.*, 1997; Cuajungco *et al.*, 2000; Beyer *et al.*, 2009; Bosomworth *et al.*, 2013), Parkinson’s disease-like dystonia (Aydemir *et al.*, 2017), amyotrophic lateral sclerosis (Kim *et al.*, 2009; Kaneko *et al.*, 2015), multiple sclerosis (Choi *et al.*, 2016), cerebral stroke (Koh *et al.*, 1996; Yoo *et al.*, 2004; Medvedeva *et al.*, 2009), and Mucolipidosis type IV (MLIV) (Cuajungco and Lees, 1997b; Eichelsdoerfer *et al.*, 2010; Cuajungco *et al.*, 2014; Smailhodzic *et al.*, 2014; Scarr *et al.*, 2016). In the case of MLIV, a neurodegenerative disease caused by the loss-of-function mutation in the transient receptor potential mucolipin-1 (TRPML1) ion channel, we found abnormal lysosomal zinc accumulation among zinc-treated MLIV patient fibroblasts and human embryonic kidney (HEK)-293 cells upon RNA interference (RNAi)-induced knock down of TRPML1 (Eichelsdoerfer *et al.*, 2010; Cuajungco *et al.*, 2014). We also observed increased tissue zinc levels in post-mortem brain of Trpml1-null mice (Eichelsdoerfer *et al.*, 2010). Our interest in human TMEM163 stemmed from these observations and our recent finding that TMEM163 interacts with TRPML1 (Cuajungco *et al.*, 2014). Rodent Tmem163, also known as synaptic vesicle 31 (SV31), was first identified from rat brain synaptosomes using proteomics analysis and was shown to bind zinc, nickel, and copper (Burre *et al.*, 2007). Tmem163 has been detected in certain types of glutamatergic and γ-aminobutyric acid (GABA)-ergic neuronal populations (Burre *et al.*, 2007; Barth *et al.*, 2011). Barth *et al*. (2007) used immunocytochemistry and subcellular fractionation studies of neuron-like PC-12 cells stably expressing rodent Tmem163 to reveal that this protein is detected mostly in the plasma membrane (PM), lysosomes, early endosomes, and other vesicular compartments especially synaptic vesicles (Barth *et al.*, 2011). Confocal microscopy studies of heterologously expressed TMEM163 in HEK-293 cells revealed that it localizes in the PM and lysosomes as well (Cuajungco *et al.*, 2014). Our analysis shows that TMEM163 transcripts are detected in many tissues, notably in the brain, lung, pancreas, kidney, ovary, and testis (Cuajungco *et al.*, 2014); however, mouse Tmem163 was reported to be exclusively expressed in the brain (Burre *et al.*, 2007).

Previously, it was shown that purified rodent Tmem163 protein reconstituted in liposomes forms a dimer and transports zinc in a proton-dependent manner (Waberer *et al.*, 2017). It was also shown that Tmem163’s ability to bind and transport zinc becomes inactivated upon alanine substitution of two species-conserved aspartate residues at position 123 and 127 (i.e. Tmem163-D123A/D127A). Thus, the data indicate that these double aspartate residues are zinc binding sites, but that they do not affect Tmem163 dimerization (Waberer *et al.*, 2017). Note, however, that this particular study did not address the question of whether the protein serves to transport zinc in or out of living cells. Here, we show for the first time that human TMEM163 is a zinc efflux transporter. Various zinc flux assays using wild-type and variant proteins confirmed the efflux transport activity of TMEM163. Overall, these observations indicate that TMEM163 should be referred to as ZNT11 (SLC30A11) being a new member of the mammalian ZNT (SLC30) family of zinc efflux transporters that are responsible for maintaining zinc homeostasis in specific cell types.

## MATERIALS AND METHODS

### Bioinformatics analysis

We performed bioinformatics analysis (www.expasy.org) of TMEM163 using TMPred (https://embnet.vital-it.ch/software/TMPRED_form.html) and TMRPres2d (Spyropoulos *et al.*, 2004) to determine and visualize its predicted transmembrane domain and topology. To identify amino acid residues subject to post-translational modification (PTM) within TMEM163, we searched the UniProt database (https://www.uniprot.org/uniprot/Q8TC26) and found several PTM entries from the PhosphoSitePlus® (https://www.phosphosite.org/proteinAction?id=24760) website as potential phosphorylation sites. We performed a Clustal W sequence alignment using Lasergene MegAlign v. 15 (DNAStar, Madison, WI) to show amino acid sequence conservation among several vertebrate species. To show amino acid sequence similarity of specific protein regions or domains between TMEM163 and select CDF family of zinc transporters, we used the Multiple Alignment using Fourier Transform (MAFFT) online resource, and consequently allowed us to determine the phylogenetic relationship between TMEM163 and certain CDF family of proteins (https://mafft.cbrc.jp/alignment/server/).

To address the previous report that mouse Tmem163 is exclusively expressed in the brain, we obtained the existing Mouse ENCODE transcriptome data (Accession no. PRJNA66167) from the National Center for Biotechnology Information (NCBI) website (https://www.ncbi.nlm.nih.gov/gene/?term=Mus+musculus+Tmem163#gene-expression). We rearranged the tissue positions to highlight the presence of Tmem163 transcripts present in other tissues.

We used the single nucleotide polymorphism (SNP) database from NCBI (dbSNP; www.ncbi.nlm.nih.gov/snp) to identify specific nucleotide sequence variations within the *TMEM163* gene that result in non-synonymous amino acid substitutions (**Supplemental Table S1**). Based on the predicted secondary structure information we obtained from several online resources (e.g. NCBI, UniProt), we selected four non-synonymous SNPs that we surmised to disrupt TMEM163’s native structure and function, namely: Serine to Arginine at position 61 (S61R), Serine to Cysteine at position 95 (S95C), Serine to Proline (S193P), and Glutamate to Lysine at position 286 (E286K). These SNPs were PCR-cloned using the primer sets outlined in **Supplemental Table S2**. Note that the PhosphoSitePlus® website mentioned above shows that Serine at position 61 is a phosphorylation site (**Supplemental Table S3**).

### Cell culture

Human embryonic kidney (HEK)-293 and HeLa cells were purchased from American Type Culture Collection (ATCC; Manassas, VA). We cultured HeLa and HEK-293 cells in Dulbecco’s Modified Eagle’s Media (DMEM; 4.50 g/L glucose, 0.58 g/L L-glutamine, 0.11 g/L sodium pyruvate; Corning) supplemented with 10% fetal bovine serum (FBS; Thermo Scientific, Waltham, MA), but without antibiotics. The cells were maintained in a humidified 37°C incubator supplied with 5% CO_2_.

### Creation of stable cell lines

We created HeLa cells that stably over-express an empty pmCherry vector control (Takara Bio, Mountain View, CA), a monomeric red fluorescent protein variant (ex = 587 nm, em = 610 nm) (Shaner *et al.*, 2004) and a TMEM163 construct with mCherry fused at the C-terminus region (Cuajungco *et al.*, 2014). Endogenous *TMEM163* transcripts are not detected in HeLa cells (www.proteinatlas.org). The pmCherry N1 vector contains a *Neomycin* resistance gene driven by the SV40 promoter. The TMEM163-mCherry and empty pmCherry vectors were linearized using the *Not I* restriction site and column-purified. The linearized vectors were transfected with TurboFect into a 70% confluent HeLa cells cultured in a 6-well plate. Every two to three days, the DMEM media containing the G418 antibiotic (0.9 mg/mL) was replaced. After two weeks, most of the untransfected cells were dead and the islands of transfected cells exhibiting red fluorescence viewed under an inverted IX-71 Olympus fluorescent microscope (Olympus America, Center Valley, PA) were isolated using cloning cylinders held in place by 1% agarose. The cloning cylinders acted as small wells from which the clonal group could be trypsinized. The dissociated colony cells were re-seeded into a 24-well culture plate. The process was repeated several times to guarantee that untransfected cells were eliminated. We then used fluorescence microscopy to corroborate that each cell line was stably expressing TMEM163 or the plasmid. As additional verification, total RNA was extracted from the stable cell lines using Trizol (Thermo Scientific), and the purified RNA samples were reverse-transcribed to generate cDNAs. The veracity of stable gene expression was established by a standard polymerase chain reaction technique using specific primer sets that anneal only to the mCherry cDNA sequence or the mCherry-tagged *TMEM163* cDNA sequence.

### RNA interference (RNAi)

We used a short hairpin RNA (shRNA, 29-mer; OriGene Technologies, Rockville, MD) expression vector that effectively down-regulates TMEM163 expression. Expression validation of shRNA knockdown using PCR and Western blot can be found in the last supplemental figure of our previous report ((Eichelsdoerfer *et al.*, 2010; Cuajungco *et al.*, 2014). For this study, we seeded 15,000 HEK-293 cells per well in PDL-treated 96-well culture plate the day before the experiments. Cells in quadruplicate wells were transfected with 2 μg DNA of either TMEM163-KD (knock-down) shRNA or TMEM163-OE (over-expression) construct using TurboFect lipid reagent (Thermo Scientific). Twenty-four hours post-transfection, the cells were washed three times with phosphate-buffered saline (PBS), exposed to low levels of exogenous zinc chloride (ZnCl_2_, 100 nM) overnight, and washed three times with PBS before the spectrofluorometric assay.

### Recombinant DNA constructs and site-directed mutagenesis (SDM) technique

We used a TMEM163-mCherry expression construct (Eichelsdoerfer *et al.*, 2010; Cuajungco *et al.*, 2014) to change a specific amino acid sequence corresponding to the non-synonymous TMEM163 SNPs used in the study (**Supplemental Table S2**). Using the online In-Fusion™ HD (IF) cloning system primer design website (Takara Bio USA, Mountain View, CA), we made primer sets for TMEM163 SNP, ZIP1 (a kind gift of Dr. David J. Eide, University of Wisconsin), and the TMEM163 double aspartate to alanine substitution mutation located at positions 124 and 128 (D124A/D128A) (**Supplemental Table S3**). We used the human ZIP1 influx transporter as a treatment control since it has been reported to localize within the PM of cells (Gaither and Eide, 2001), which is similar to the TMEM163 protein localization (Cuajungco *et al.*, 2014). Note that the double aspartate to alanine mutations were previously reported to decrease zinc binding and inactivate protein function in rodent Tmem163 at positions 123 and 127 (Tmem163-D123A/D127A) (Waberer *et al.*, 2017). Following SDM of the D124A/D128A and SNP constructs, we verified the DNA sequence integrity of all expression vectors by commercial sequencing (Retrogen, San Diego, CA) and analyzed the clone sequences using Lasergene SeqMan v. 15 (DNAStar, Madison, WI).

### Spectrofluorometric assays using Fluozin-3 and Newport Green fluorescent dyes

#### TMEM163 RNAi expression

Following RNAi and exposure to zinc, the cells were washed three times with PBS. The cells were incubated with 1 µM of cell membrane permeable Fluozin-3 acetoxy-methyl ester (AM) fluorescent dye (MP-FZ3; Kd = 15 nM; ex = 494 nm, em = 516 nm; Thermo Scientific). Fluozin-3 is a well-established tool for evaluating changes in relative zinc levels due to its specificity, high affinity, and physiological stability (Zhao *et al.*, 2008). The dye only emits fluorescence when bound to labile or chelatable zinc and the dye remains in the cytoplasm upon ester cleavage of the AM moiety (Gee *et al.*, 2002). We followed the manufacturer’s recommendation to incubate MP-FZ3 for 30 minutes inside a 37°C incubator injected with 5% CO_2_. The cells were then washed with PBS three times before performing an end-point assay for intracellular zinc fluorescence as previously described (Eichelsdoerfer *et al.*, 2010; Cuajungco *et al.*, 2014). Untransfected negative control cells were included in all experiments (n = 4 independent trials). Background fluorescence was subtracted (blank) from each treatment group to obtain the relative fluorescence unit (RFU). The data were analyzed using Prism 8.2 software (GraphPad, San Diego, CA).

#### Stable TMEM163 cell expression

We optimized the spectrofluorometric assay for cell membrane impermeable Fluozin-3 (MI-FZ3) and the low affinity, membrane permeable Newport Green dichloro-fluorescein-AM (MP-NG; Kd ~ 1 μM; ex = 505 nm, em = 535 nm; Thermo Scientific) by identifying the appropriate: (i) cell density; (ii) concentration of exogenous ZnCl_2_; (iii) concentration of MI-FZ3 or MP-NG; and (iv) buffer types for dye incubation and washing. We also supplemented both the incubation and wash buffers with 2 mM Glutamax™ (Thermo Scientific) and 1 mM sodium pyruvate to maintain cell viability during experimentation.

We used HeLa cells stably expressing TMEM163-mCherry or pmCherry empty vector control. We seeded 25,000 cells per well in a PDL-treated 96-well plate one day before the experiments. The first six wells of each row were designated as treatment group (zinc exposed), while the remaining six wells in the same row were used as untreated control group (non-exposed). We then exposed the cells to ZnCl_2_ (10 μM) and zinc pyrithione (1 μM) for 5 minutes and washed five times with HEPES buffer (15 mM HEPES, 100 mM D-glucose, and 150 mM KCl; pH 7.0) using an automated BioTek 405 TS microplate washer (BioTek Instruments, Winooski, VT). A post-washed volume of 100 μL was dispensed and the plate was transferred into the BioTek Cytation V plate reader and each well was dispensed with 100 μL HEPES buffer containing MI-FZ3 dye to a final concentration of 1 μM. We performed kinetic readings for each well every minute for 25 minutes. Average background RFU from non-zinc exposed group (transfected and non-transfected, but not pre-loaded with zinc) was subtracted from RFUs of zinc-exposed group (transfected and non-transfected). Background subtraction minimizes or prevents the confounding effects contributed by native zinc transporters present in cells such as ZNTs and ZIPs, which are both expressed in HeLa and HEK-293 cell lines (Human Protein Atlas, www.proteinatlas.org). Five-minute data point intervals were collected and analyzed using Prism 8.2 software.

To further support the MI-FZ3 results, we then performed assays using MP-NG. HeLa cells stably expressing TMEM163 and pmCherry vector control were seeded at 25,000 cells per well in a PDL-treated 96-well plate. The cells were incubated with MP-NG (5 μM) for 20 minutes inside a 37°C incubator injected with 5% CO_2_, washed two times with a standard buffer (10 mM HEPES, 135 mM NaCl, 5 mM KCl, 1 mM CaCl_2_, 1 mM MgSO_4_, and 5 mM glucose; pH 7.4), and left at room temperature for another 30 minutes to ensure complete ester cleavage of the AM moiety. The treatment group was exposed to ZnCl_2_ (100 μM) and pyrithione (10 μM) for 20 minutes inside the incubator, washed five times with standard buffer, and each well was dispensed with a final volume of 100 μL standard buffer after the last wash. A kinetic reading for zinc flux was performed every minute for 45 minutes using the BioTek Cytation V plate reader. Background intracellular MP-NG RFU levels (no zinc exposure) were subtracted from the zinc-treated RFU levels. We adapted a previously described protocol (Ohana *et al.*, 2009) to quantify changes in cytoplasmic zinc levels by dividing the RFU timepoint readings with the baseline RFU taken at time zero. Data points from 5-minute interval reads were collected and analyzed using Prism 8.2 software.

#### Transient TMEM163 cell expression

To further corroborate the stable expression data and include additional controls, we performed transient transfection of the following constructs: pmCherry empty vector, TMEM163-mCherry, ZIP1-mCherry, and TMEM163-D124A/D128A-mCherry (a similar double mutation reported to inactivate rodent Tmem163) (Waberer *et al.*, 2017). We seeded HEK-293 cells at 15,000 cells per well on a 96-well culture plate treated with PDL one day before transfection. The first six wells of each row were designated as treatment group (zinc exposed), while the remaining six wells in the same row were used as untreated control group (non-exposed). We transfected the cells with 150 ng of DNA constructs per well using TurboFect lipid reagent. A standard amount of 200 ng DNA was then used per well for the empty vector control to match the efficiency of transfection in cells treated with the expression constructs containing a specific gene. Twenty-four hours post-transfection, the culture medium was aspirated, and each well was dispensed 100 μL of HEPES buffer with zinc (treated) or without zinc (untreated). The treated cells were exposed to ZnCl_2_ (10 μM) and zinc pyrithione (1 μM) for 5 minutes in a humidified 37°C incubator supplied with 5% CO_2_. The cells were then washed three times with HEPES buffer using an automated BioTek 405 TS microplate washer to minimize or prevent cells from lifting out of the well surface, and after the final liquid aspiration, each well was dispensed with 100 μL of HEPES buffer containing MI-FZ3 dye to a final concentration of 1 μM. The plate was transferred into the BioTek Cytation V plate reader and each well was assayed (kinetic reading) for zinc flux every minute for 25 minutes. To analyze the raw data, the background RFU values (no zinc exposure) were subtracted from zinc-treated RFU data as described earlier. Five-minute data point intervals were collected and analyzed (n ≥ 5 independent trials) using Prism 8.2 software.

To show that the MI-FZ3 fluorescence signal is directly attributable to zinc, we used the metal chelator N,N,N,N-*tetrakis*-(2-pyridylmethyl)-ethylenediamine (TPEN, 10 μM) in several transient transfection experiments as described earlier by injecting the chelator half-way through the kinetic reading. TPEN is a zinc-specific, high affinity chelator (Kd = 10^−16^ M) that binds chelatable zinc *in vivo* (Cuajungco and Lees, 1996; 1998) or *in vitro* (Zalewski *et al.*, 1993; Ahn *et al.*, 1998). One-minute interval reads in quadruplicate wells were analyzed and graphed using Prism 8.2 software.

To evaluate the effects of each non-synonymous SNP on TMEM163 function, we seeded HEK-293 at 15,000 cells per well in a PDL-treated 96-well plate the day before the experiment. Similar to the above approach for MI-FZ3, the first six wells were designated as zinc-treated group, while the last six wells were used as non-zinc treated group. We used TurboFect to transfect the cells with the following non-synonymous SNP-associated constructs (150 ng DNA): TMEM163-S61R-mCherry, TMEM163-S95C-mCherry, TMEM163-S193P-mCherry, and TMEM163-E286K-mCherry. Twenty-four hours post transfection, the cells were pre-loaded with ZnCl_2_ (10 μM) and zinc pyrithione (1 μM) for 20 minutes, washed twice with HEPES buffer, and transferred to the plate reader for kinetic reading after injection of the MI-FZ3 dye (1 μM). A kinetic reading for zinc flux was performed every minute for 25 minutes using the BioTek Cytation V plate reader and the background RFU values were subtracted from the zinc-treated RFU values. Data points every 5 minutes were collected for analysis (n ≥ 4 independent trials) and graphed using Prism 8.2 software.

### Radionuclide zinc-65 uptake assay

We purchased radionuclide zinc-65 (^65^Zn, 1 µCi; Perkin Elmer, Waltham, MA) and followed the zinc flux assay protocol published by the Eide laboratory (Gaither and Eide, 2001) with minor modifications to suit our experimental needs. Each treatment was done in duplicate wells, and a parallel plate for normalization using total protein levels was included in each experiment. HeLa cells stably expressing TMEM163-mCherry or pmCherry vector were seeded at 200,000 cells per well in a 12-well culture plate treated with PDL Twenty-four hours later, the cells were washed three times with uptake buffer (15 mM HEPES, 100 mM glucose and 150 mM KCl; pH 7.0). Different concentrations of ^65^Zn (40 µM – 1.25 µM) were prepared in the uptake buffer. The cells were incubated with 250 µl of the various ^65^Zn solutions for 15 minutes inside a 37°C incubator injected with 5% CO_2_. Following ^65^Zn exposure, the plate was placed on ice and 1 ml of cold stop buffer solution (1 mM EDTA in uptake buffer) was added in each well to chelate extracellular ^65^Zn in the milieu. EDTA is a membrane impermeable chelator that binds zinc (Kd = 10^−16^ M) (Cuajungco and Lees, 1998; Frederickson *et al.*, 2002). The cells were washed three times with cold stop buffer. After the last wash, the buffer was aspirated and the cells were lysed with 500 µl PBS containing 1% Triton-X-100. The ^65^Zn-treated cell lysates were assayed for radioactivity in counts per minute (cpm) using a gamma counter. The information provided by the manufacturer regarding the specific activity of ^65^Zn allowed us to convert the cpm data to pmol per minute.

A parallel untreated plate was lysed for subsequent use in total protein assay. The lysis buffer used consists 50 mM Tris-HCl, 150 mM NaCl, 1% Nonidet P-40, 1 mM EDTA, 0.25% sodium deoxycholate, 0.1% SDS, 1mM PMSF and 1X protease inhibitor cocktail (Sigma-Aldrich, St. Louis, MO).The relative total protein concentration of lysates from TMEM163-mCherry and pmCherry-expressing cells were quantified (in milligrams) using the Pierce Bicinchoninic acid (BCA) assay kit according to the manufacturer’s protocol (Thermo Fisher Scientific). The intracellular ^65^Zn levels were then normalized by the total protein concentration. We used the Prism 8.2 software to graph the results and determine statistical significance between the TMEM163 and control data (n = 3 independent trials).

### Immunocytochemistry (ICC) and fluorescence microscopy analysis

HEK-293 cells were seeded in sterile coverslips at approximately 30% confluency one day before transfection of the following expression constructs: TMEM163 wild-type (TMEM163-WT), TMEM163-D124A/D128A, TMEM163-S61R, TMEM163-S95C, TMEM163-S193P, TMEM163-E286K, ZIP1, and pmCherry vector control. The mammalian expression constructs used in the study either contained a fluorescent (mCherry; Takara Bio) protein tag or peptide tag (Myc-DDK; OriGene Technologies).

For mCherry-tagged constructs, we checked for transfection efficiency and the health of treated cells at 24 hours post-transfection before processing the cells for microscopy. The cells were fixed with 4% paraformaldehyde for 15 minutes, and washed three times with 1X PBS. Fluorescence imaging was done using an inverted Olympus IX-71 fluorescence microscope. Monochromatic RGB images of cells were captured using the CellSens software version 1.18 (Olympus). We then used Adobe Photoshop CC 2017 (Adobe Systems, San Jose, CA) to show the images in red color as represented by the mCherry fluorescence.

For Myc-DDK-tagged constructs, the cells were fixed with 4% paraformaldehyde for 15 minutes, washed three times with PBT1 buffer (1X PBS, 1% Triton-X 100, 0.1% BSA, 5% heat-inactivated goat serum) and incubated with primary antibody (1:1000 anti-DDK monoclonal 4C5; OriGene Technologies) in PBT1 at 4°C overnight. The cells were washed with PBT2 buffer (1X PBS, 1% Triton-X 100, 0.1% BSA) three times, and then incubated with secondary antibody (1:500 anti-mouse Alexa Fluor-488; ex = 490 nm, em = 525 nm; Thermo Scientific) in PBT2 for 2 hours at 4°C. The cells were then washed two times with 1X PBS. Each coverslip was mounted on a slide, treated with Prolong Gold anti-fade reagent and the edge of the coverslip was sealed with clear nail polish. Monochrome images were taken using an upright Olympus BX51 fluorescence microscope and processed using Adobe Photoshop CC 2017 to change the monochrome images to green color as represented by Alexa-488 fluorescence.

### Cell viability assay

To investigate whether the SNPs and the TMEM163-D124A/D128A constructs affect cell health, we seeded 15,000 HEK-293 cells in a PDL-treated 96-well plate. Cells in quadruplicate wells were transfected with TMEM163-WT, TMEM163-D124A/D128A, TMEM163-S61R, TMEM163-S95C, TMEM163-S193P, TMEM163-E286K, and pmCherry empty vector control. We then assessed cell viability of treated and control cells 48 hours post-transfection using the CellTiter Glo 2.0 luciferase assay kit (Promega, Madison, WI) according to the manufacturer’s protocol. The data were normalized using un-transfected control cells and represented as percentage of control, and statistical significance was analyzed using Prism 8.2 software (n = 5 independent trials).

### Cell surface biotinylation (CSB) and Western blot (WB)

HeLa cells were seeded at 1.5 × 10^6^ cells in a 10-cm petri dish coated with PDL. We transfected the cells with Myc-DDK peptide tag plasmids (4 µg) using TurboFect lipid reagent as follows: TMEM163-WT, TMEM163-S61R, TMEM163-S95C, TMEM163-S193P, TMEM163-E286K and TMEM163-D124A/D128A. Twenty four hours post-transfection, the cells were washed two times with 10 ml of ice-cold PBS-CM (1X PBS, 0.5 mM CaCl_2_, 1 mM MgCl_2_; pH 8.0) and incubated with 1 ml of freshly prepared Sulfo-NHS-LC-LC-Biotin solution (Thermo Scientific; 0.5 mg/ml in PBS-CM) for 30 min at 4°C. The cells were washed twice with 10 ml of PBS-CM containing 0.1% BSA to terminate and quench any unbound Sulfo-NHS-LC-LC-Biotin. The cells were washed again with PBS (pH 7.4) and then lysed for 1 hour at 4°C with 1 ml of lysis buffer (1X protease inhibitor cocktail, 1mM PMSF, 150 mM NaCl, 5 mM EDTA, 1% Nonidet P-40, 50 mM Tris; pH 7.5). The lysates were centrifuged at 14,000 rpm for 10 min at 4°C. The relative total protein concentration of the lysates was quantified (in milligrams) using the BCA assay kit according to the manufacturer’s protocol (Thermo Fisher Scientific).The neutravidin beads (150 µl; Thermo Scientific) were equilibrated in lysis buffer, added to each protein sample (normalized to 680 µg total protein), and incubated overnight at 4°C. The samples were washed five times with lysis buffer, incubated in 60 µl 2X Laemmli sample buffer for 30 min at 37°C, and eluted by low speed centrifugation. The samples were loaded in 4-12% gradient Bis-Tris NuPAGE (Thermo Fisher Scientific) SDS-PAGE gel and were immunoblotted with primary anti-DDK monoclonal mouse antibody (1:5000) (Origene Technologies). The Western blot was viewed using secondary anti-mouse IR-Dye 800CW (1:10000) (LI-COR Biosciences). The blots were then scanned using the Odyssey SA™ infrared imaging system (LI-COR Biosciences).

### Statistical analysis

All numerical data were graphed and analyzed for statistical significance using Prism 8.2 software (GraphPad, La Jolla, CA). We used Student’s *t*-test (two-tailed) to determine statistical significance of data from various zinc flux assays using stable cells expressing TMEM163 and pmCherry vector control. The Km and Vmax values of efflux transport activity were obtained using the non-linear curve fit analysis on Prism. Data obtained from other spectrofluorometric experiments were analyzed using one-way analysis of variance (ANOVA) with repeated measures and Tukey’s multiple comparisons *post-hoc* test. The significance level was set at *p* < 0.05. Unless specified, all data evaluated for statistical significance are represented as mean ± SEM with at least three independent trials.

## RESULTS

### TMEM163 is a zinc effluxer and is a member of the CDF family of proteins

TMEM163 has six predicted transmembrane (TM) domains and is highly conserved among the vertebrate species we analyzed (**Supplemental Figure S1**). RNAi of TMEM163 in HEK-293 cells produced a substantial increase of intracellular MP-FZ3 fluorescence, while over-expression of TMEM163 displayed an opposite effect (**Supplemental Figure S2**). This result confirmed and was consistent with our previous report (Cuajungco *et al.*, 2014). Thus, we had an *a priori* information that TMEM163 may be transporting zinc out of living cells, but needed more evidence to substantiate this observation.

For this study, we used two different types of human cell lines, two heterologous expression systems (i.e. stable and transient), and three zinc flux assays to show the specificity and reproducibility of our observations that TMEM163 is an efflux transporter. To first show that TMEM163 effluxes zinc, we used HeLa cells stably expressing TMEM163 and analyzed zinc flux upon exogenous zinc exposure using two zinc-specific fluorescence dyes, MI-FZ3 or MP-NG. For MI-FZ3 assay, we found a substantial increase in extracellular fluorescence among TMEM163-expressing cells in comparison to the pmCherry control (Fig.1A). For MP-NG, a substantial reduction of intracellular MP-NG fluorescence was observed from cells expressing TMEM163 when compared to the pmCherry control (Fig. 1B). To provide additional evidence confirming our fluorescence data that TMEM163 is a zinc efflux transporter, we exposed HeLa cells stably expressing TMEM163-mCherry and mCherry with radionuclide ^65^Zn. We found that TMEM163-expressing cells significantly reduced intracellular ^65^Zn levels when compared with control cells (Fig.1C).

**Figure 1.**
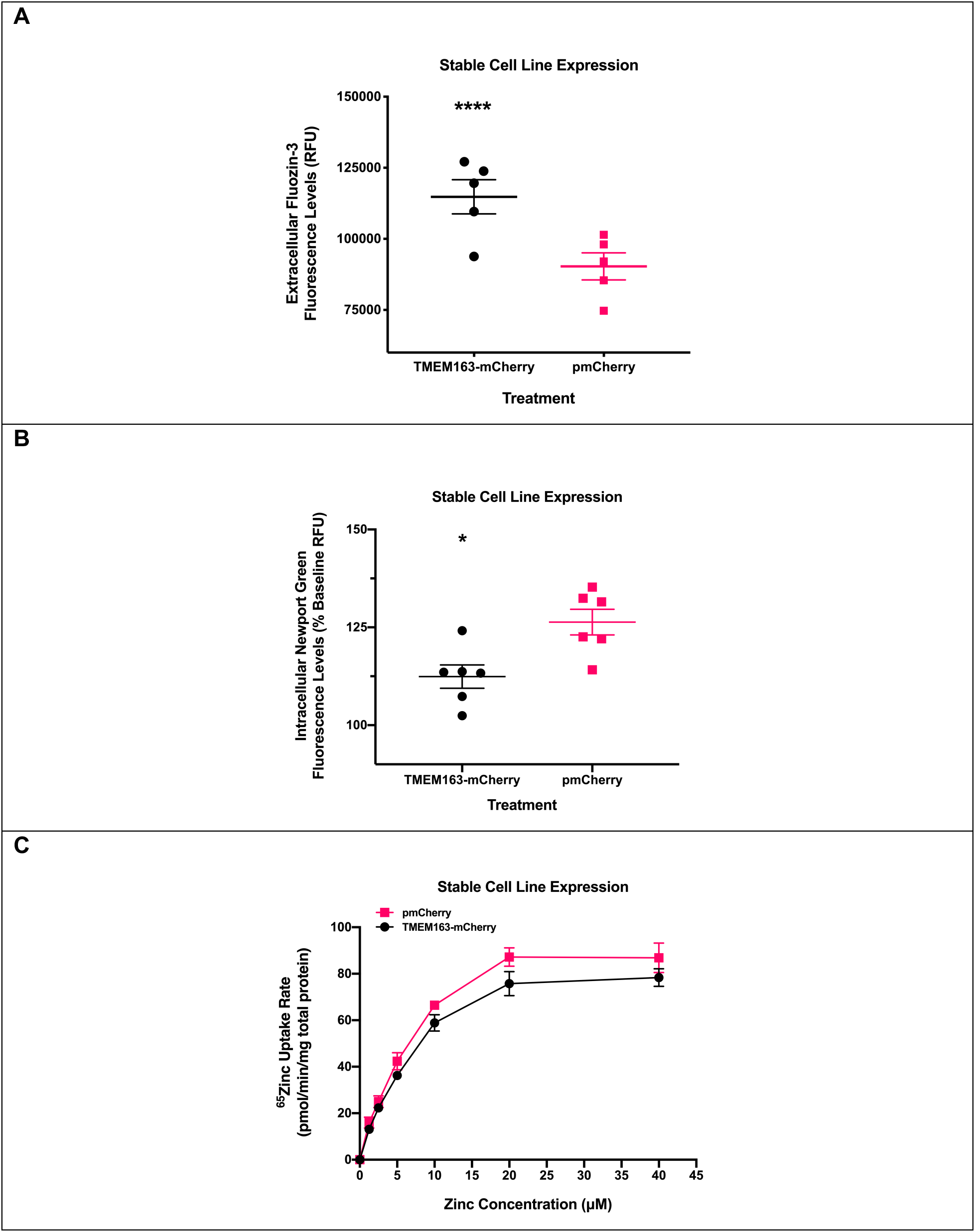
Functional expression of TMEM163 stably expressed in HeLa cells. **A**) Cell membrane impermeant Fluozin-3 (MI-FZ3) fluorescence analysis revealed a significant increase in extracellular RFU levels in the milieu of cells expressing TMEM163-mCherry upon exposure to zinc. Data are represented as mean ± SEM (\*\*\*\**p* < 0.0001, Student’s *t*-test, unpaired, two-tailed; n = 5 independent trials). **B**) Cell membrane permeant Newport Green (MP-NG) fluorescence analysis revealed a significant decrease in intracellular RFU levels of cells expressing TMEM163-mCherry upon exposure to zinc when compared with cells expressing the pmCherry vector control. Data are represented as mean ± SEM of each kinetic time point analyzed (\**p* = 0.01; Student’s *t*-test, unpaired, two-tailed; n = 6 independent trials). **C**) Concentration dependence and saturable uptake of ^65^Zn by TMEM163-expressing and mCherry-expressing cells. The presence of TMEM163 exhibited significant reduction of intracellular ^65^Zn levels (*p* = 0.01; Student’s *t*-test, paired, two-tailed; n = 3 independent trials).

We investigated the possibility that TMEM163 is indeed a member of the CDF family of zinc efflux transporters by performing multiple sequence alignment (MSA) and phylogenetic tree analyses using MAFFT. Comparison of the full-length TMEM163 amino acid sequence with those of non-mammalian CDF proteins YiiP, CzcD, and ZitB, as well as mammalian CDF proteins, ZNT1 through ZNT10, suggested that TMEM163 is a new member of the CDF family (Figure 2). We confirmed that TMEM163 shared conserved zinc-binding sites consisting of two aspartic acid residues within TM2 (Barth *et al.*, 2011). In addition, we identified aspartic residues located within TM5 and TM6 that could be potential zinc-binding sites, which are conserved or shared by several distinct CDF members. Lastly, the phylogenetic relationship among CDF proteins hinted that TMEM163 is more related to ZNT9 in comparison to other CDF family members that were analyzed.

**Figure 2.**
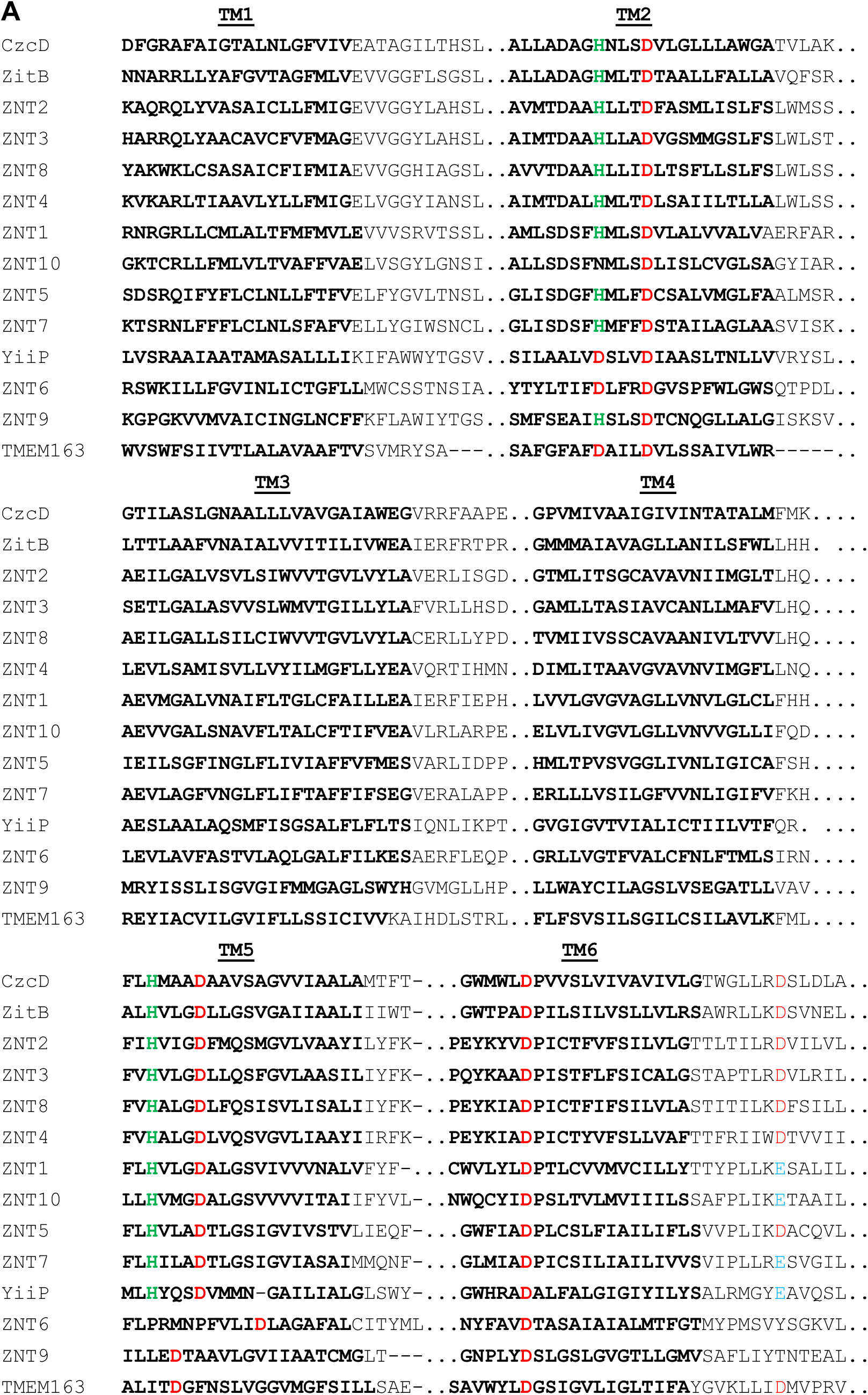
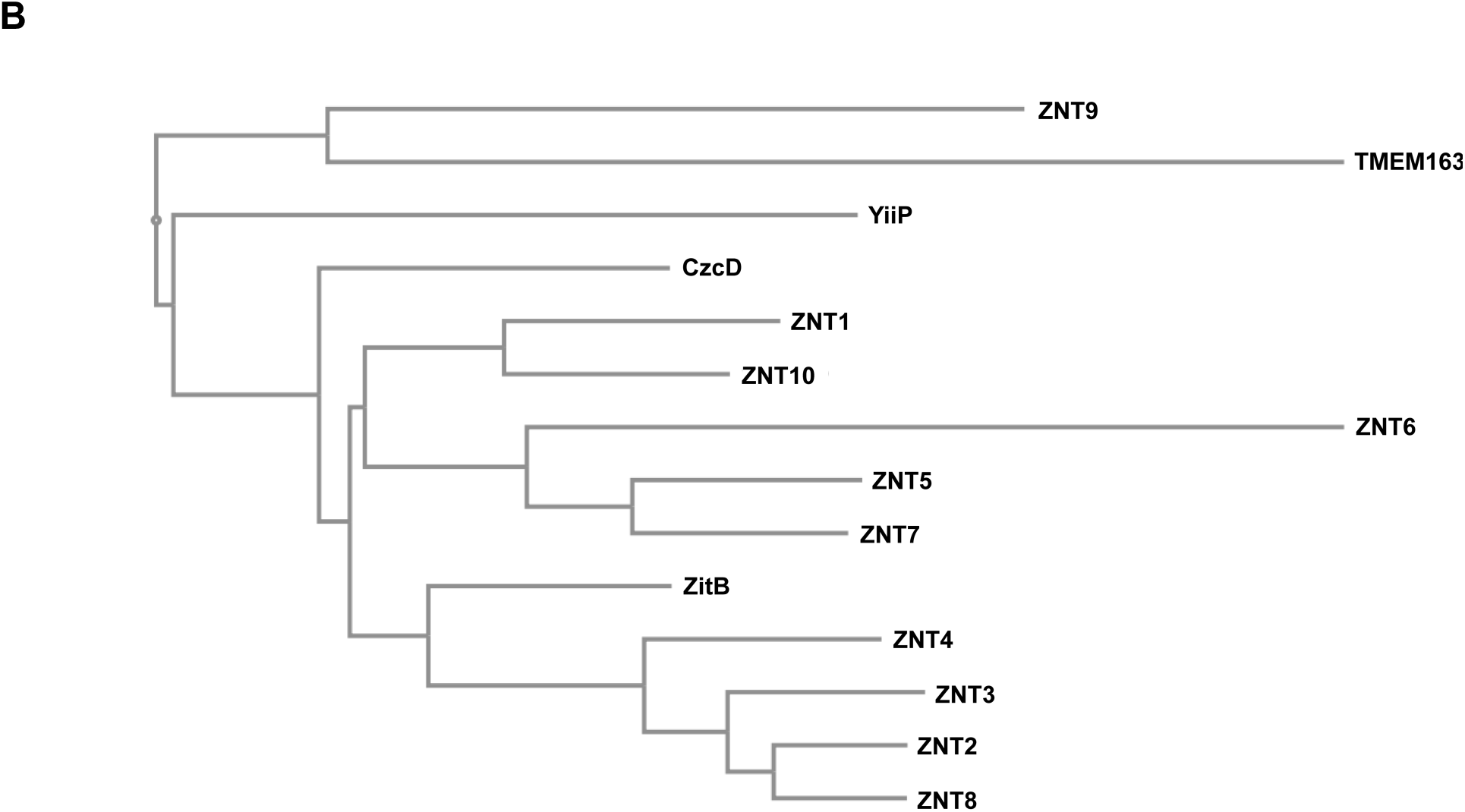
Multiple sequence alignment of TMEM163 and select CDF family members. **A**) Clustal alignment of TMEM163 and CDF amino acid sequences showing predicted transmembrane domains 1 through 6 (TM1-TM6; *bold text*). The alignment was done using the MAFFT website (https://mafft.cbrc.jp/alignment/server/) for the following full-length protein sequences obtained from NCBI: CzcD (YP_002515680.1), TMEM163 (NP_112185.1), YiiP (NP_462942), ZitB (WP_000951292), ZNT1 (NP_067017.2), ZNT2 (NP_001004434.1), ZNT3 (NP_003450.2), ZNT4 (NP_037441.2), ZNT5 (NP_075053.2), ZNT6 (NP_001180442.1), ZNT7 (NP_598003.2), ZNT8 (NP_776250.2), ZNT9 (NP_006336.3), and ZNT10 (NP_061183.2). For simplicity, only TM domains (*bold text*) and consensus or conserved amino acid residues predicted to bind zinc ions are shown. Histidine (*green*), Aspartic acid (*red*), and Glutamic acid (*blue*). **B**) Phylogram of the evolutionary relationship among specific CDF family members and TMEM163 protein. The phylogenetic tree diagram was generated using the Java-based Archeopteryx visualization and analysis of phylogenetic trees via the MAFFT web interface.

### Loss-of-function mutation negatively affects TMEM163 function but not membrane localization

We looked at the effects of transient heterologous expression of TMEM163 in HEK-293 cells. We used pmCherry empty vector as a negative control and ZIP1 as a treatment control because it is a zinc influx transporter that is present in the PM just like TMEM163. We also included the double aspartate to alanine substitutions, TMEM163-D124A/128A, to verify that this variant will be inactive, since this clone is equivalent to rodent Tmem163-D123A/D127A and was previously reported as an inactive protein (Waberer *et al.*, 2017). We observed a discernable increase of MI-FZ3 fluorescence in the extracellular milieu of TMEM163-expressing cells, but not among cells expressing pmCherry empty vector control, ZIP1, and TMEM163-D124A/D128A (Fig. 3A). Statistical analysis using ANOVA with repeated measures, and *post-hoc* analysis using Dunnett’s multiple comparisons test showed significant differences between TMEM163 and: i) TMEM163-D124A/D128A (*p* < 0.0001); ii) ZIP1 (*p* < 0.0001); and iii) pmCherry (*p* = 0.002). To demonstrate that the MI-FZ3 fluorescence signal is contributed by zinc extrusion in the extracellular environment, we performed additional experiments in which we injected TPEN during the kinetic measurement. TPEN is a high affinity, cell membrane permeant chelator that binds tightly to zinc (Cuajungco and Lees, 1996; 1998). We found that TPEN completely abrogated the MI-FZ3 fluorescence signal (**Supplemental Fig. S3**).

**Figure 3.**
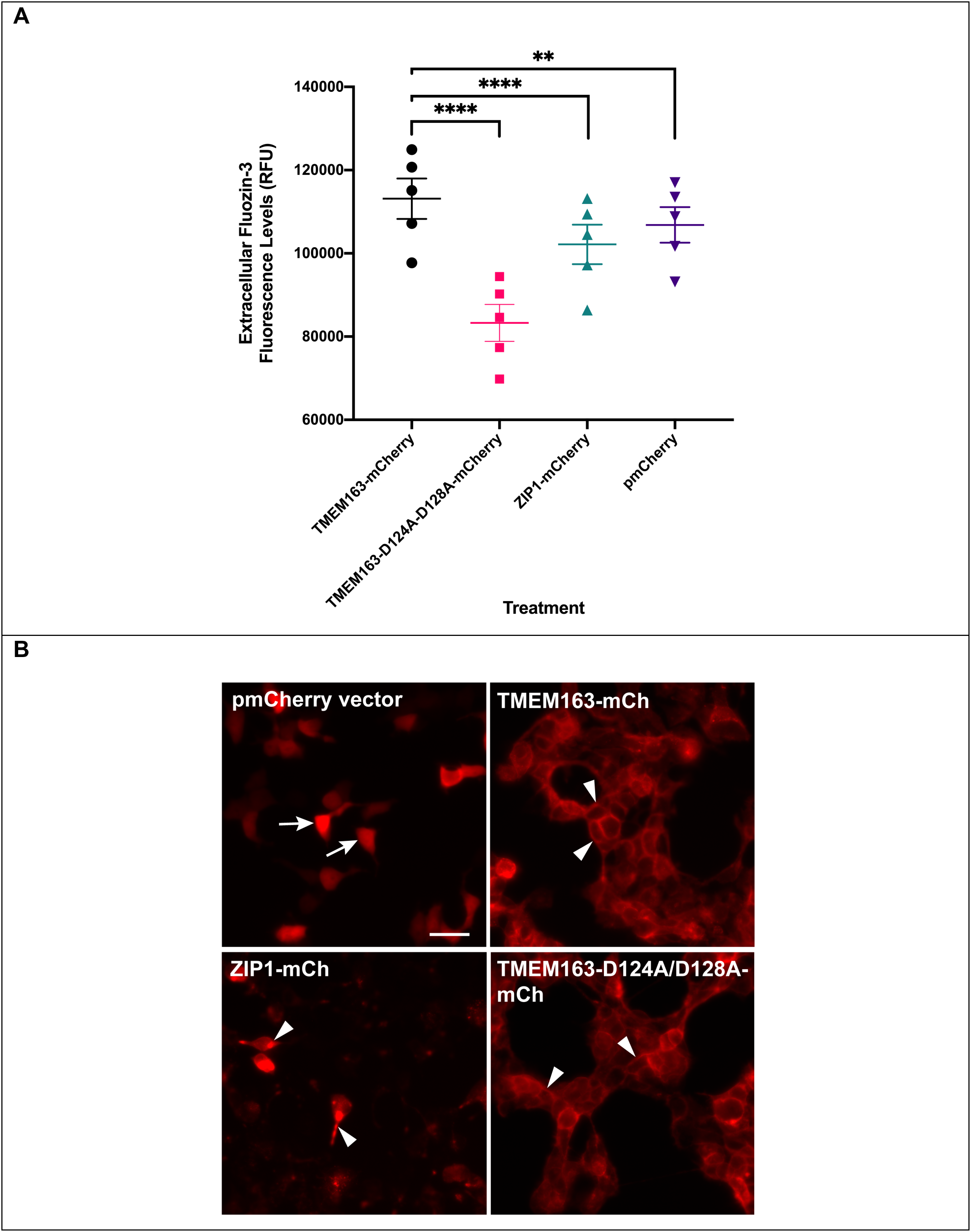
Spectrofluorometric assay and fluorescence microscopy of HEK-293 cells transiently expressing wild-type TMEM163, TMEM163-D124A/128A, ZIP1, and pmCherry empty vector control. **A**) Cell membrane impermeant Fluozin-3 fluorescence analysis reveals a significant increase in extracellular fluorescence upon expression of TMEM163 protein when compared with controls. Data are represented as mean ± SEM of each kinetic time point analyzed (\*\**p* = 0.002, \*\*\*\**p* < 0.0001; Tukey’s multiple comparisons *post-hoc* test; n ≥ 5 independent trials). **B**) Representative fluorescent micrographs showing subcellular distribution of pmCherry (mCh) empty vector, TMEM163-mCherry, ZIP1-mCherry, and TMEM163-D124A/D128A-mCherry. TMEM163 and TMEM163-D124A/D128A localize within the plasma membrane and membrane compartments (*arrowhead*). ZIP1 is also detected within the plasma membrane and intracellular membrane compartment (*arrowhead*). The pmCherry vector exclusively localizes in the cytoplasm (*arrow*). Scale bar: 50 µm.

We wanted to ensure that the marked difference in zinc efflux activity between TMEM163-WT and TMEM163-D124A/D128A was not caused by either transfection or cell expression issues. We therefore took fluorescent images of the cells for evidence of transfection before performing the assays. We also performed fluorescence microscopy to determine the expression profiles of all four constructs. Our findings revealed that TMEM163 is localized within the plasma membrane (PM) and intracellular membrane compartments (MC) (Fig. 3B), which further confirmed our previous report (Cuajungco *et al.*, 2014). Interestingly, the subcellular localization of TMEM163-D124A/D128A appeared to be similar to the wild-type protein where it is detected in both the PM and MC. ZIP1 expression was detected in the PM and more prominently within MC, which is consistent with previous reports (Gaither and Eide, 2001; Wang *et al.*, 2004). As expected, the pmCherry empty vector was mainly observed in the cytoplasm.

### Missense mutations in conserved amino acid residues negatively affects the zinc efflux function of TMEM163

Based on the apparent lack of transport activity by the D124A/D128A variant, we hypothesized that a single point mutation within certain conserved amino acid residues could potentially disrupt, if not also abolish, the normal efflux function of TMEM163. To avoid simply selecting amino acids based on information from prediction software tools, we strategically looked at specific non-synonymous SNPs that have been identified within the *TMEM163* gene using NCBI’s dbSNP database (**Supplemental Table S1**). We picked four non-synonymous SNPs of serine (i.e. S61R, S95C, and S193P) and glutamic acid (i.e. E286K) residues due to their species conservation, position within the predicted secondary structure, and possible targets for post-translational modification (**Supplemental Figure S4**). While amino acid substitution within serine (phosphorylation site) or glutamic acid (negatively charged) residue may negatively impact protein structure and function, our *in silico* analysis suggested that only S61 is a potential PTM site (**Supplemental Table S3**). We transiently expressed the non-synonymous SNP constructs in HEK-293 cells along with wild-type TMEM163 and pmCherry empty vector control. For this set of experiments, we incubated the cells with zinc and pyrithione for an extra 15 minutes than the previous TMEM163 transfection experiments to gauge whether the amino acid substitution could affect cell health upon longer zinc exposure. We found that the cells expressing the SNP constructs showed significant reductions in extracellular MI-FZ3 fluorescence signal when compared with wild-type TMEM163 (Fig. 4A). ANOVA with repeated measures and *post-hoc* analysis using Dunnett’s multiple comparisons test showed significant differences between wild-type TMEM163 and: i) S61R (*p* = 0.001); ii) S95C (*p* = 0.04); S193P (*p* = 0.0004); and iii) E286K (*p* = 0.0005). The MI-FZ3 fluorescence signals coming from cells expressing S61R, S95C, and S193P constructs were remarkably similar to that of cells expressing the pmCherry empty vector control. Note, however, that the cells expressing the E286K construct displayed the lowest RFU levels(Fig. 4A) and resembled the data trend obtained for the D124A/D128A protein variant (Fig. 3A).

**Figure 4.**
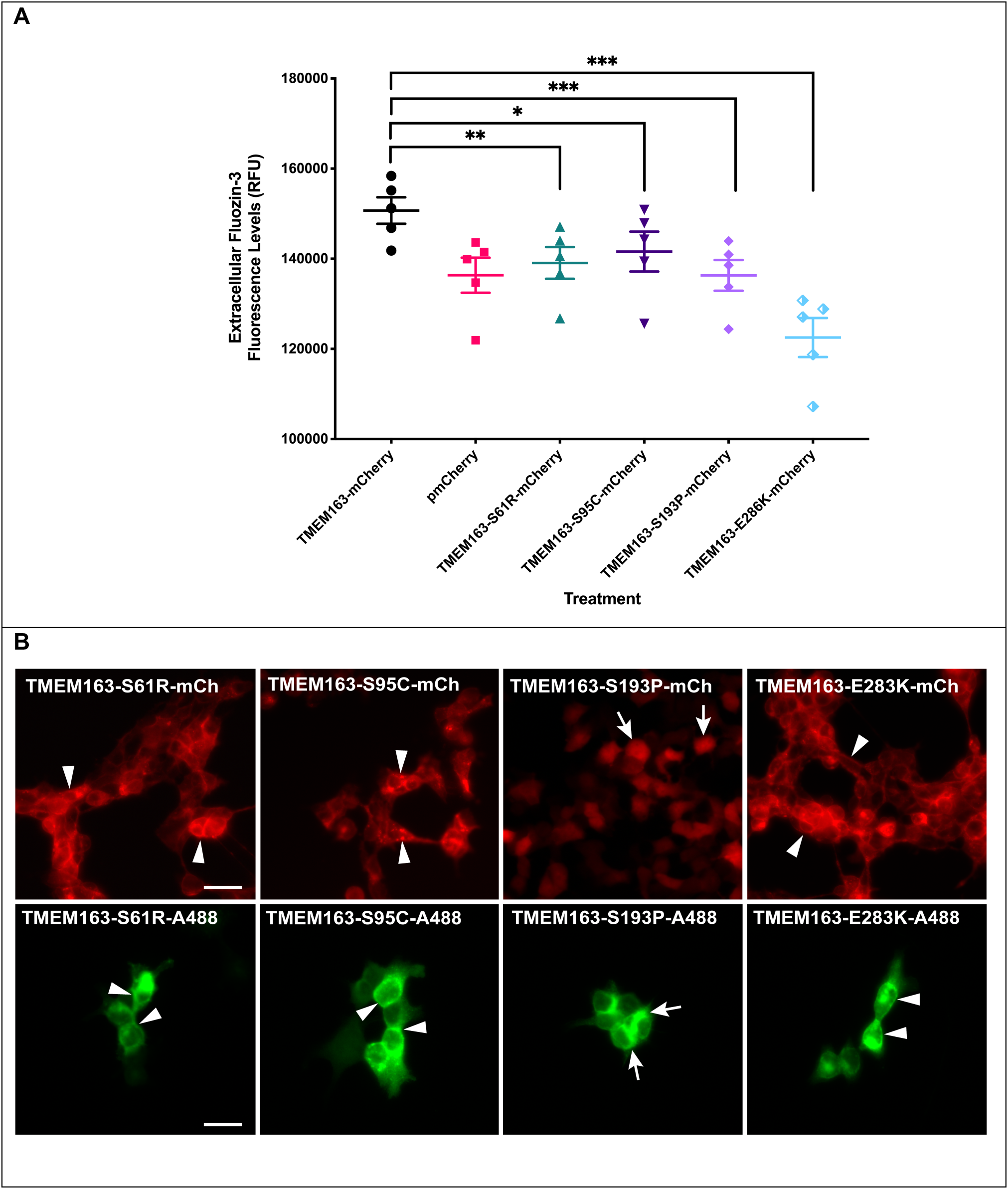
Spectrofluorometric assay and fluorescence microscopy of HEK-293 cells transiently expressing wild-type TMEM163 and non-synonymous SNPs: S61R, S95C, S193P, and E286K. **A**) Cell membrane impermeant Fluozin-3 fluorescence analysis reveals a significant increase in extracellular fluorescence upon expression of TMEM163 protein. Non-synonymous SNPs negatively affect the efflux function of TMEM163, particularly the E286K clone. Data are represented as mean ± SEM of each kinetic time point analyzed (\**p* = 0.04, \*\**p* = 0.004, \*\*\**p* = 0.0004; Tukey’s multiple comparisons *post-hoc* test; n ≥ 4 independent trials). **B**) Representative fluorescent micrographs of expression constructs with mCherry (mCh) fluorescent protein tag (*top panel*) and Myc-DDK peptide tag probed with Alexa Fluor-488 (A488) (*bottom panel*). TMEM163-S61R, TMEM163-S95C, and TMEM163-E286K exhibit localization within the plasma membrane and intracellular membrane compartment (*arrowhead*), while TMEM163-S193P shows mostly cytoplasmic and compartmental localization (*arrow*). Scale bar: 50 µm.

### Most of the TMEM163 variant alleles localize to the plasma membrane and do not affect cell viability

Similar to our earlier experimental approach, we captured fluorescence images of the cells expressing the SNPs to determine transfection efficiency and expression profile before we performed each MI-FZ3 assay. Fluorescence microscopy analyses of SNP constructs tagged with mCherry fluorescent protein or Myc-DDK peptide showed analogous expression pattern to that of the wild-type TMEM163 protein despite differences in their efflux activity (Fig. 4B). It is worth noting that the S193P-mCherry expression profile appeared exclusively within the cytoplasm, which paralleled the empty pmCherry vector control expression in cells. On the other hand, the expression of S193P-Myc-DDK was mainly observed within MC, but a PM localization was also detected. Evaluation of cell surface protein levels using CSB and WB demonstrated that all SNPs, including the D124A/D128A variant were present within the PM; however, the level of S193P protein variant was markedly reduced relative to the normalized samples of other SNPs (**Supplemental Fig. S5**). The CSB and WB data also revealed that S61 might be a PTM target (i.e. phosphoserine) based on the protein band migration pattern of S61R protein variant, which was below the expected bands for wild-type TMEM163, D124A/D128A, and other SNP protein variants.

To provide additional confirmation that the decline of zinc efflux activity among non-synonymous SNPs and D124A/D128A constructs was not caused by diminished cell health, we performed cell viability assays and found no indication of cytotoxicity in these cells (**Supplemental Fig. S6**). Although the cells expressing the S95C construct exhibited relatively lower viability, the values were not significant when compared to wild-type TMEM163 and other treatments. Overall, these results further support the evidence that TMEM163 is a zinc effluxer. Figure 5 summarizes the main findings of the study.

**Figure 5.**
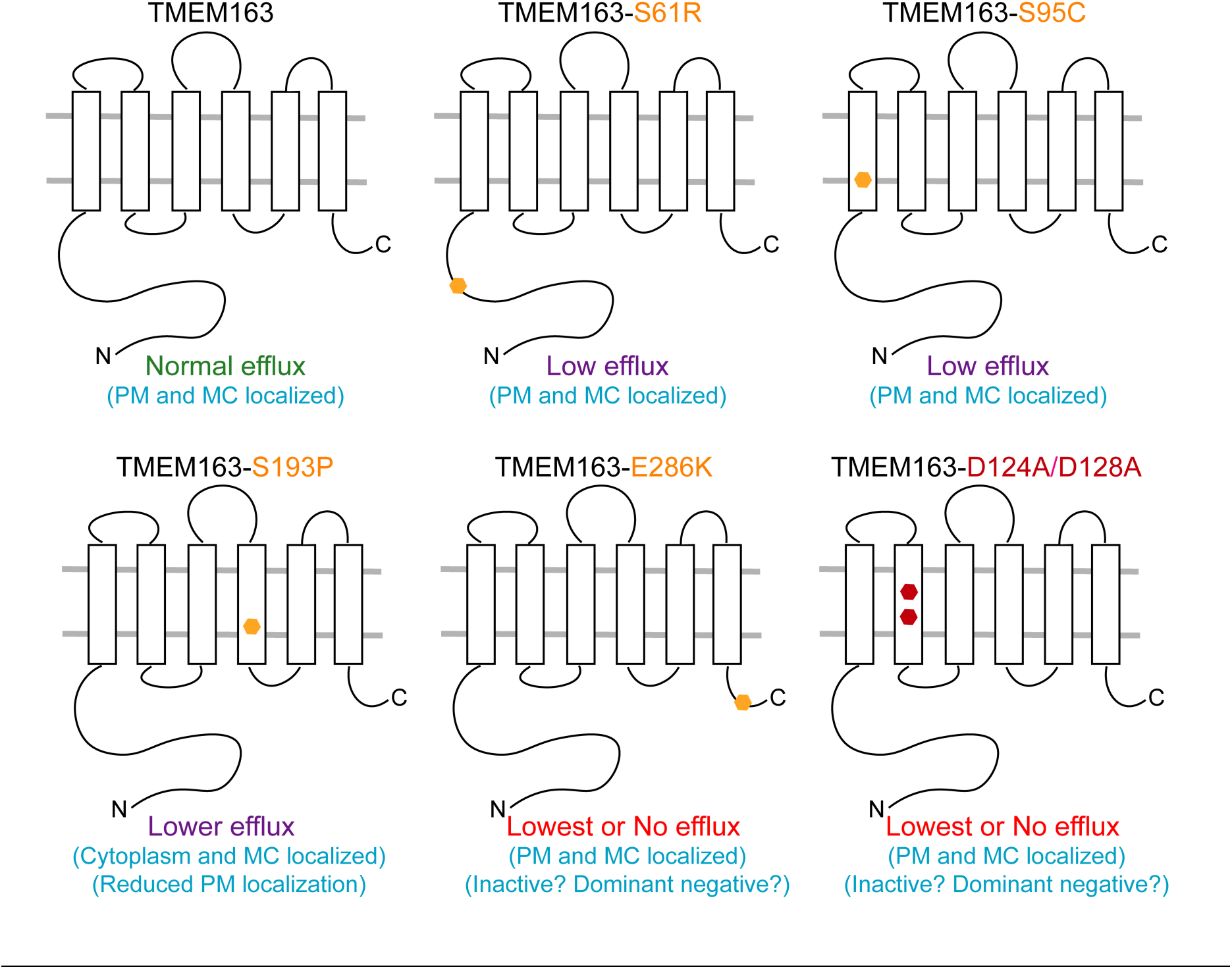
Schematic diagram summarizing the main observations on the zinc efflux function of TMEM163. Heterologous expression of the wild-type TMEM163 results in cytoplasmic zinc extrusion in the presence of high zinc levels inside the cells. Expression of the non-synonymous SNPs S61R, S95C, and S193P produces marked reduction of efflux activity when compared with wild-type TMEM163. S61R showed altered protein band migration pattern on WB, suggesting that S61 is a PTM target. S193P is mostly expressed in the cytoplasm and MC as evidenced by CSB and WB, while the other variants are found mostly in the PM and MC. The E286K protein variant has the lowest efflux activity in comparison with other SNPs and wild-type TMEM163. Similar to E283K, the D124A/D128A protein variant has comparably the lowest efflux activity relative to the wild-type TMEM163 and other SNPs. PM, plasma membrane; MC, membrane compartment.

### Mouse Tmem163 transcript expression is not exclusive to the brain

To address the previous report that the expression of mouse *Tmem163* transcripts is only detected in the brain tissue (Burre *et al.*, 2007), we searched for publicly available RNA sequencing (RNA-seq) data on *Tmem163* expression. We found the Mouse ENCODE transcriptome data on NCBI, which clearly illustrated that *Tmem163* transcripts are present in several tissues of adult and embryonic mice, in addition to the cerebral cortex and cerebellum. Specifically, the RNA-seq data show relatively high expression levels of *Tmem163* mRNA in heart, lung, and spleen (**Supplemental Figure S7**), suggesting that Tmem163 may also serve an important role in these tissues.

## DISCUSSION

Our investigations on the direction of zinc flux facilitated by TMEM163 protein present within the PM ultimately revealed that it serves to transport zinc out of the cell. Our interests on human TMEM163 protein stemmed from our previous discovery that it is an interaction partner of the Transient Receptor Potential Mucolipin-1 (TRPML1) protein (Cuajungco *et al.*, 2014). Recently, it was shown that rodent Tmem163 interacts with itself and forms a functional dimer (Waberer *et al.*, 2017). Despite being predicted as an efflux transporter, earlier reports on rodent Tmem163 (Barth *et al.*, 2011; Waberer *et al.*, 2017) and our work on human TMEM163 (Cuajungco and Kiselyov, 2017) have suggested that the protein might be operating as an influx transporter. A case in point, it was reported that PC12 cells that heterologously express rodent Tmem163 displayed a noticeable rise in MP-FZ3 fluorescence within the cytoplasm upon exposure to zinc (Barth *et al.*, 2011). This report suggested that Tmem163 transports extracellular zinc inside the cells (Barth *et al.*, 2011). Furthermore, the same group showed that purified Tmem163 in liposomes mediate luminal zinc accumulation as evidenced by Fluozin-1 fluorescent dye when these lipid nanodiscs were exposed to exogenous zinc (Waberer *et al.*, 2017). In our previous preliminary study, TMEM163-expressing HEK-293 cells acutely exposed with zinc produced a discernible up-regulation of *MT-1A* transcripts after twenty-four hours when compared with control cells (Cuajungco and Kiselyov, 2017), which suggests that TMEM163 might be an influxer. In contrast, when we knocked down TMEM163 expression in HEK-293 cells using RNAi, the fluorescence intensity of MP-FZ3 dye inside the cells remained considerably higher than in control cells (Cuajungco *et al.*, 2014). After carefully re-analyzing these contradictory observations and upon performing additional experiments, we hypothesized that TMEM163 might bring zinc out of the cells in response to high cytosolic zinc levels. We reasoned that endogenous zinc transporters (e.g. ZNTs and ZIPs) present in cells confounded our previous interpretation of increased MT-1A transcript levels upon TMEM163 expression. It may be that over-expressing TMEM163 increases zinc extrusion from cells, and this effect induces a compensatory exogenous zinc uptake through native zinc influxers, which in turn, induces the expression of *MT-1A* transcripts. In line with this reasoning, transcripts of several zinc influx *SLC39* genes (e.g. ZIP1, ZIP3, ZIP6-ZIP10, ZIP13, and ZIP14) are detected in HEK-293 and HeLa cells (www.proteinatlas.org). Meanwhile, transcripts of certain *SLC30* genes are also detected in HEK-293 cells (e.g. ZNT5 and ZNT6) and HeLa cells (e.g. ZNT1, ZNT5, and ZNT6) (www.proteinatlas.org). Waberer and colleagues (2017) circumvented the potential confounding effects of these native zinc influx and efflux proteins by using artificial liposomes. In their report, they purified rodent Tmem163 proteins and inserted them into lipid nanodiscs. This approach allowed them to show that Tmem163 binds to and transports zinc as evidenced by an increase of cell membrane permeant Fluozin-1 fluorescence inside the liposomes (Waberer *et al.*, 2017). While this elegant cell-free system provided evidence for Tmem163-mediated zinc transport, it did not show whether the protein is an influxer or effluxer in living cells. Despite challenges in utilizing human cell lines in the current study, we successfully showed that TMEM163 extrudes zinc out of cells upon intracellular zinc elevation. Coupled with the MSA and phylogenetic data, it is reasonable to propose that TMEM163 should now be formally called ZNT11 protein as a new member of the human ZNT (SLC30) family of zinc efflux transporters. Meanwhile, future investigations to determine the Michaelis-Menten constant (Km) of TMEM163-mediated efflux using radionuclide ^65^Zn in a cell-free system are needed, in order to compare its kinetics with that of rodent Tmem163, whereby a Km value of 44.3 μM was calculated from Fluozin-1 fluorescence data (Waberer *et al.*, 2017).

The dramatic negative response of TMEM163-D124A/D128A variant on zinc efflux in comparison to ZIP1 and pmCherry vector control was unexpected, because if this variant is a non-functional protein as has been reported for rodent Tmem163-D123A/D127A (Waberer *et al.*, 2017), then we would expect the magnitude of fluorescence signal for this variant to be similar to that of pmCherry empty vector control. Our data show that the depressed efflux activity of the D124A/D128A variant is not due to mis-localization and/or cell toxicity as demonstrated by fluorescence imaging and viability assays, respectively. Since rodent Tmem163 has been reported to form a dimer, it is conceivable that the TMEM163-D124A/D128A monomer acts in a dominant negative manner upon dimerization with native TMEM163 monomers present in HEK-293 cells. We also cannot rule out the prospect that TMEM163-D124A/D128A may interact and induce a dominant negative effect on native zinc transporters present in HEK-293 cells (e.g. ZNTs), since various zinc efflux transporters have been reported to form heterodimers (Fukunaka *et al.*, 2009; Salazar *et al.*, 2009; Lasry *et al.*, 2014; Golan *et al.*, 2015). It would be interesting to find out in future work if the potential dimerization of D124A/D128A variant with native TMEM163 or with other native zinc transporters could be responsible for the apparent dominant negative effect on zinc efflux. Nevertheless, the data obtained in this study is further substantiation that it TMEM163 a zinc efflux transporter.

For experiments studying the consequence of non-synonymous SNPs in HEK-293 cells, we found that the amino acid substitution in three serine residues (S61R, S95C, and S193P) or in one glutamic acid residue (E286K) completely reduced TMEM163’s zinc efflux activity. Interestingly, the CSB and WB results support the possibility that S61 is a PTM site. It is conceivable that the presence of arginine in the S61R variant disrupted protein modification as shown by the protein band migration pattern on the Western blot when compared with wild-type TMEM163 and other protein variants. It would be interesting in future research to determine other PTM sites predicted to be located within the N-terminus of TMEM163, and to identify specific enzymes that are involved in the modification of this region. Meanwhile, the apparent mis-localization of S193P regardless of its C-terminus tag (mCherry fluorescent protein or dual Myc-DDK peptide), suggests that the proline substitution within the predicted TM4 of TMEM163 could have kinked this domain’s alpha-helical structure – an effect that is reminiscent of the TRPML3 alanine to proline substitution seen in the varitint-waddler mouse (Grimm *et al.*, 2007; Cuajungco and Samie, 2008). Interestingly, the mis-localization of S193P appears to be more severe in the presence of the mCherry fluorescent protein tag, possibly due to the size of mCherry, which has a molecular mass of 26.7 kDa (Shaner *et al.*, 2004). The reduction of PM localization also explains why the S193P protein variant has a lower efflux activity compared to the wild-type protein. On another note, it is not clear why the zinc efflux activity of E286K is dramatically reduced, which is virtually comparable to D124A/D128A variant. It is plausible that the change from a negatively charged glutamate to a positively charged lysine residue altered the protein’s structure, which then created a dominant negative mutant like that of the D124A/D128A protein variant. Future studies on TMEM163 structure and function could explain the probable dominant negative effect of site-specific mutations within the protein. Nevertheless, the reduction of zinc efflux activity observed from these non-synonymous SNPs provided additional support that TMEM163 is an effluxer.

Meanwhile, our analysis of the mouse Tmem163 transcript expression suggests that, like its human counterpart (Cuajungco *et al.*, 2014), this gene is detected in various tissues and is not exclusively expressed in the mouse brain as previously reported (Burre *et al.*, 2007). Whether the observed mouse transcripts in the brain and other tissues are translated into a functional protein needs further investigation and is beyond the scope of the current study.

## CONCLUSIONS

Our findings reveal that TMEM163 may play a crucial role in cellular zinc homeostasis by extruding cytoplasmic zinc ions to the extracellular environment. Noteworthy is that TMEM163 is unique in that it has a short C-terminus region, whereas the human ZNT family of efflux transporters have typically long C-terminus regions. Despite such structural difference, both functional similarity and evolutionary relationship among the CDF proteins and TMEM163 indicate that it should now be referred to as ZNT11 protein and its gene named as *SLC30A11*. Finally, we found that non-synonymous SNPs within the *TMEM163* (*SLC30A11*) gene may alter protein structure and negatively affect its function. Therefore, additional studies on the importance of TMEM163 (ZNT11) protein and its contributions to normal or pathological cellular states are highly warranted.

## Abbreviations

AM: Acetoxy-methyl ester
CDF: Cation diffusion facilitator
CSB: Cell surface biotinylation
CT: carboxyl terminus
EDTA: Ethylenediamine tetraacetic acid
FZ3: Fluozin-3 fluorescent dye
KD: Knockdown
MC: Membrane compartment
MI: membrane impermeable or impermeant
MP: membrane permeable or permeant
NCBI: National Center for Biotechnology Information
NG: Newport Green fluorescent dye
NT: amino terminus
OE: Over-expressed
PM: Plasma membrane
PTM: Post-translational modification
RFU: Relative fluorescence unit
RNAi: RNA interference
SDM: Site-directed mutagenesis
shRNA: short-hairpin RNA
SLC: Solute carrier
SNP: Single nucleotide polymorphism
SV31: Synaptic vesicle 31 protein
TM: Transmembrane domain
TPEN: N,N,N,N-*tetrakis*-(2-pyridylmethyl)-ethylenediamine
WB: Western blot
ZIP: Zrt- and Irt-like protein
ZNT: Zinc efflux transporter

## ACKNOWLEDGMENTS

We are very grateful to Theodros Kidane, Lauren Rosas, Cathleen Nguyen, Quinlan Cantrell, and Joshua Silva for their technical support. We also thank Dr. Sean Murray (CSU Northridge) for reading and critiquing the manuscript.

## AUTHOR CONTRIBUTIONS

MPC conceived and designed the study, analyzed and interpreted data, wrote the manuscript, and project administration. VBS performed experiments, analyzed data, and edited the manuscript. SA performed experiments including assay optimization, analyzed data, and edited the manuscript. AE performed experiments and edited the manuscript. All authors read and approved the final version of the manuscript.

