## Supplemental Data for "Transmembrane 163 (TMEM163) protein effluxes zinc"

**Table S1.** The National Center for Biotechnology Information (NCBI)'s Single Nucleotide Polymorphism database (*dbSNP*) was used study specific nucleotide sequence variations within the *TMEM163* gene. We identified SNPs that produce amino acid substitution when compared with the wild-type (WT) TMEM163 protein sequence. The information obtained was then used to generate our In-Fusion site-directed mutagenesis primers.

| dbSNP Cluster ID | mRNA Position | Function | dbSNP Allele | Amino acid Residue | Codon Position | Amino acid Position |
|---|---|---|---|---|---|---|
| *rs531340951* | 249 | Missense | A | Arginine | 3 | 61 |
| | | Contig reference | C | Serine | 3 | 61 |
| *rs202199940* | 350 | Missense | G | Cysteine | 2 | 95 |
| | | Contig reference | C | Serine | 2 | 95 |
| *rs113424259* | 643 | Missense | C | Proline | 1 | 193 |
| | | Contig reference | T | Serine | 1 | 193 |
| *rs371768391* | 922 | Missense | A | Lysine | 1 | 286 |
| | | Contig reference | G | Glutamate | 1 | 286 |

**Table S2.** Tabulated list of In-Fusion primers used in the study. The primer set to clone ZIP1 is included among primer sets used for site-directed mutagenesis (SDM) primers. The list shows SDM of specific nucleotides corresponding to the double aspartate to alanine mutation and non-synonymous SNPs located within the N- and C-termini, as well as transmembrane domain (TMD)-1 and TMD4. The primers were designed using the online In-Fusion cloning primer design tools from Takara Bio USA (Mountain View CA) website. Primers were commercially synthesized by Integrated DNA Technologies (Coralville IA). The In-Fusion PCR cloning was performed according to the manufacturer's recommendations.

| Primer Name | Primer Sequence (5' → 3') |
|---|---|
| *TMEM163-S61R (NT*)* | Fwd: CCAGTTCAGAGACGGGCTGGAGGACCGA<br>Rev: CCGTCTCTGAACTGGCCGCTCTCGCT |
| *TMEM163-S95C (TM1*)* | Fwd: CTGGTTCTACATCATTGTCACCCTGGCCC<br>Rev: ATGATGTAGAACCAGGACACCCACAATGC |
| *TMEM163-S193P (TM4*)* | Fwd: CAGTGTCCCCATTTTAAGTGGGATTCTTTGCAGC<br>Rev: AAAATGGGGACACTGAACAGGAAATCGTC |
| *TMEM163-E286K (CT*)* | Fwd: TCACTACAAGATGTTTGAGCGAATTCTGCAGTCG<br>Rev: AACATCTTGTAGTGACGTGTCTGCCTCACC |
| *TMEM163-D124A/D128A* | Fwd: TGCTGCCATCCTGGCTGTCCTGTCATCGGCGATTGTC<br>Rev: GCCAGGATGGCAGCAAATGCAAACCCAAAAGCAGAGG |
| *ZIP1* | Fwd: CTCGAGCTCAAGCTTACCATGGGGCCCTGGGGAGAGCCA<br>Rev: GACCGGTGGATCCCGGATTTGGATGAAGAGCAGGCC |

*Predicted regions: NT, N-terminus; CT, C-terminus; TM, transmembrane domain

**Table S3.** Predicted post-translational modification (PTM) within TMEM163 (UnitProt Q8TC26) using PhosphoSitePlus® (https://www.phosphosite.org/proteinAction?id=24760). Note that the phosphoserine residue at position 61 (*red text*) was identified to be a known variant with non-synonymous single nucleotide polymorphism (i.e. S61R) that was used in the current study.

| | Post-translational Modification | | | | |
|---|---|---|---|---|---|
| **AA Position** | **Human** | | **Mouse** | | **Rat** |
| *7* | Isoleucine | | Phosphoserine | | Phosphoserine |
| *11* | Phosphoserine | | Phosphoserine | | Phosphoserine |
| *12* | Phosphoserine | | Proline | | Proline |
| *16* | Phosphothreonine | | Glycine | | Glycine |
| *22* | Mono-methylarginine | *23* | Arginine | *23* | Arginine |
| *27* | Proline | *28* | Phosphoserine | *28* | Serine |
| *28* | Alanine | *29* | Phosphothreonine | *29* | Threonine |
| *34* | Alanine | *35* | Phosphoserine | *35* | Asparagine |
| *37* | Phosphoserine | | — | | — |
| *41* | Arginine | *40* | Phosphoserine | *40* | Serine |
| *42* | Glutamic acid | *41* | Phosphoserine | *41* | Serine |
| *46* | Leucine | *45* | Phosphoserine | *45* | Proline |
| *55* | Phosphoserine | *54* | Phosphoserine | *54* | Serine |
| *57* | Phosphoserine | *56* | Phosphoserine | *56* | Phosphoserine |
| *61* | Phosphoserine | *60* | Phosphoserine | *60* | Phosphoserine |
| *72* | Serine | *71* | Phosphoserine | *71* | Serine |
| *73* | Serine | *72* | Phosphoserine | *72* | Serine |
| *77* | Ubiquityllysine | *76* | Lysine | *76* | Lysine |

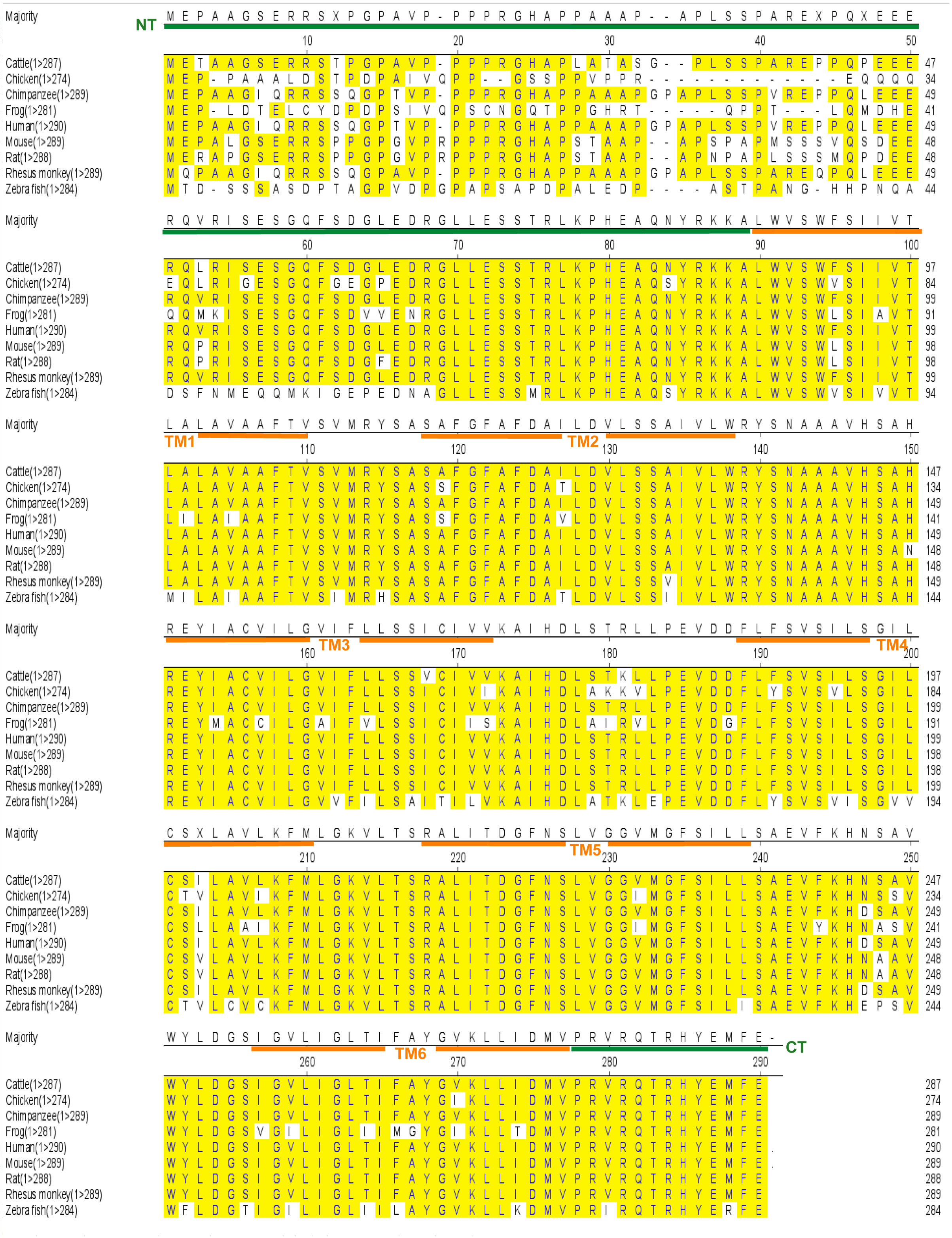

**Figure S1**. **Multiple species alignment of TMEM163 protein using Clustal W.** TMEM163 has six predicted transmembrane (TM) domain shown by the *orange* solid line, while the N-terminus (NT) and C-terminus (CT) are demarcated by the *green* solid line. Shaded amino acids (*yellow*) indicate species conservation. The illustration was generated using Lasergene MegAlign v. 15.

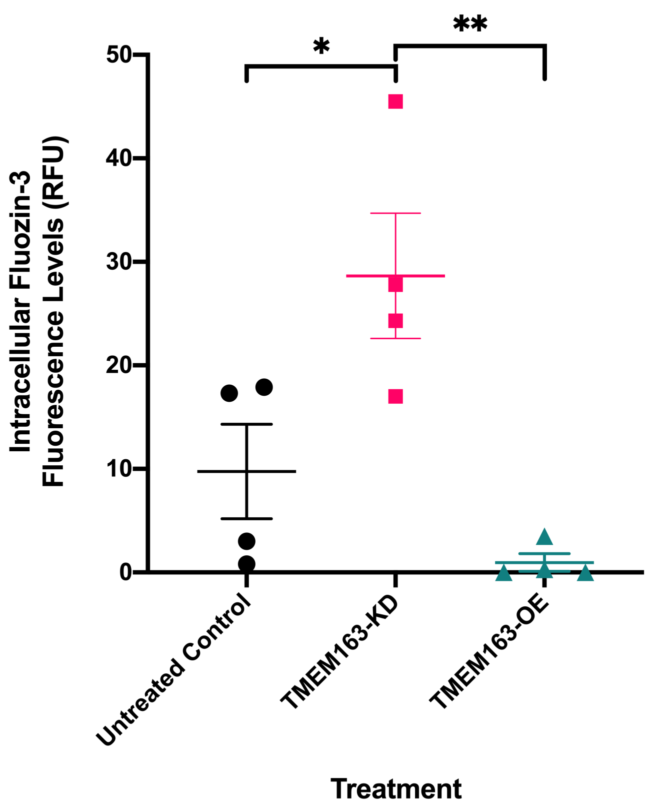


**Figure S2**. **RNAi of endogenous TMEM163 protein increases cytoplasmic zinc levels.** Cultured HEK-293 cells were incubated with membrane-permeable FluoZin-3 dye (500 nM) upon exposure to exogenous zinc ($ZnCl_2$, 100 nM) for 24 h following transient transfection of either TMEM163-shRNA construct to knockdown (KD) or TMEM163-mCherry construct to overexpress (OE) the protein. Intracellular Fluozin-3 dye fluorescence levels increased in cells with TMEM163-KD, but not in TMEM163-OE and untreated control cells, which confirmed our previous report (Cuajungco *et al.*, 2014). Data are represented as mean ± SEM of each experiment (*$p < 0.05$, **$p < 0.01$; Student's *t*-test, paired, n = 4 independent trials). Validation data of the TMEM163 shRNA using both PCR and Western blot techniques are available as supplemental material from our previous report (Cuajungco *et al.*, 2014).

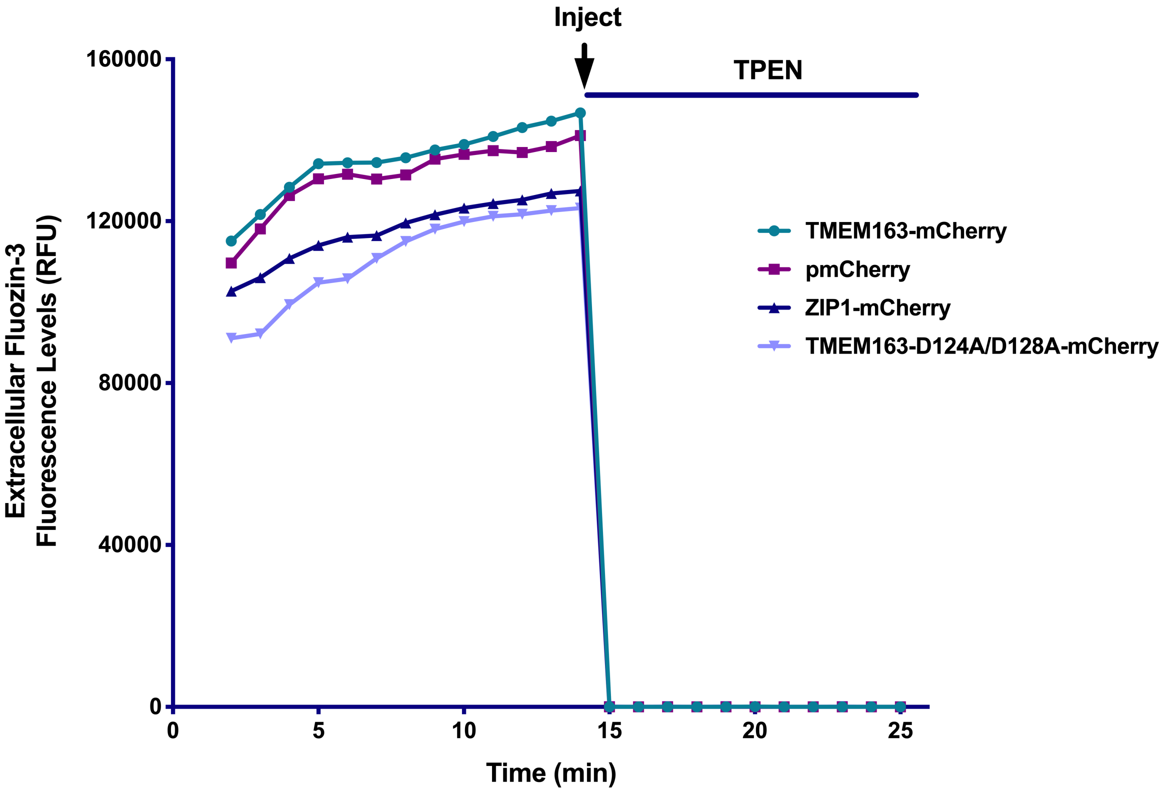


**Figure S3**. **The extracellular Fluozin-3 fluorescence signals are abrogated upon TPEN treatment.** Spectrofluorometric analysis of HeLa cells transiently expressing TMEM163-mCherry, ZIP1-mCherry, TMEM163-D124A/D128A-mCherry, and pmCherry empty vector following exposure to zinc. The cells were assayed using cell membrane impermeant Fluozin-3 dye (in quadruplicate wells). The fluorescent signals disappeared upon injection of TPEN (a high affinity zinc chelator), which indicate that the fluorescence signals are due to the presence of zinc.

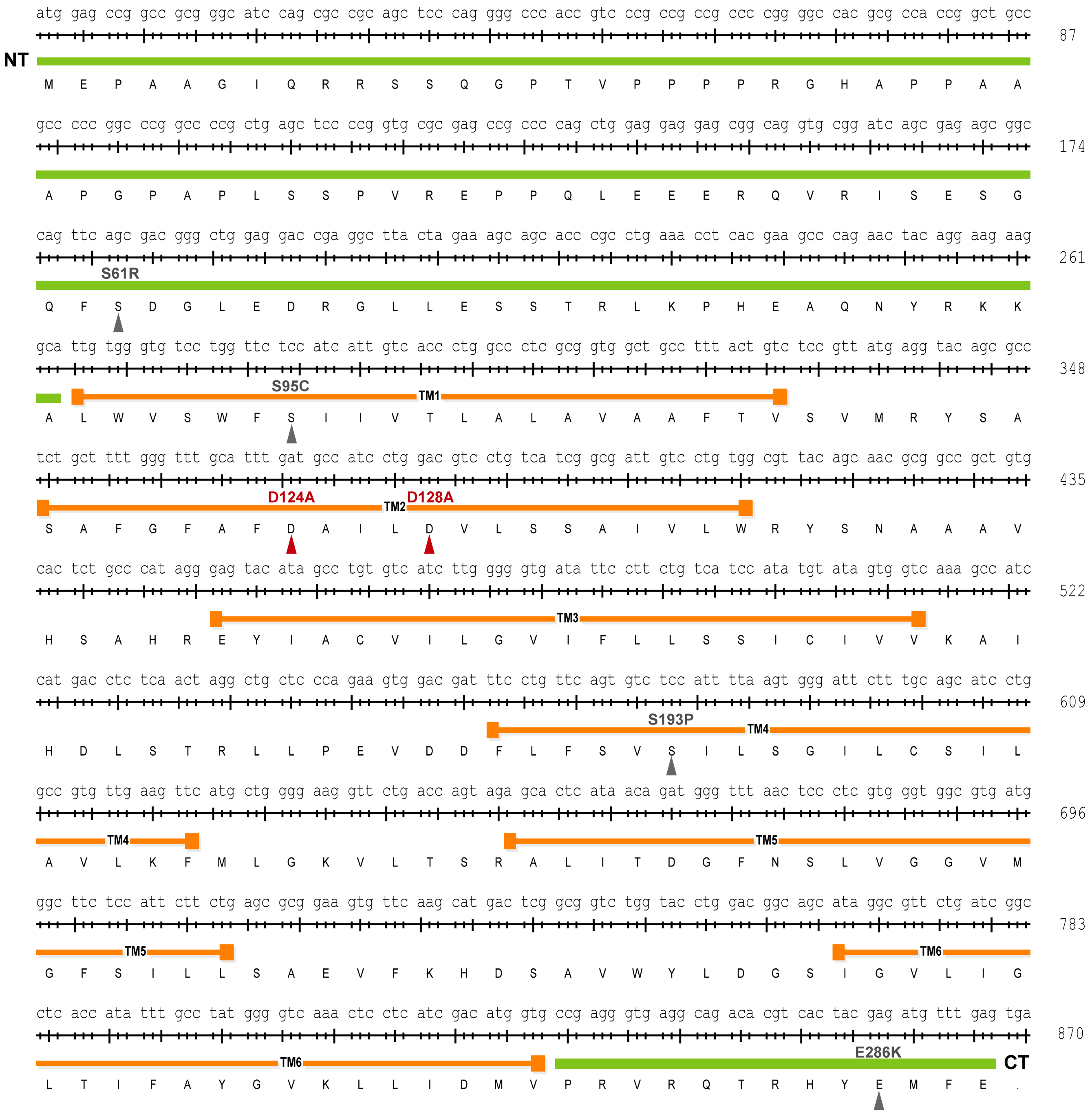


**Figure S4. Schematic diagram of *TMEM163* open-reading frame illustrating the location of amino acid residues associated with non-synonymous SNPs and the known loss-of-function double alanine to aspartic acid substitutions.** The map shows the location of D124A/D128 double mutations (*red text and symbol*) and non-synonymous SNPs S61R, S95C, S193P and E286K (*gray text and symbol*). The map was generated using Lasergene SeqBuilder v. 15.

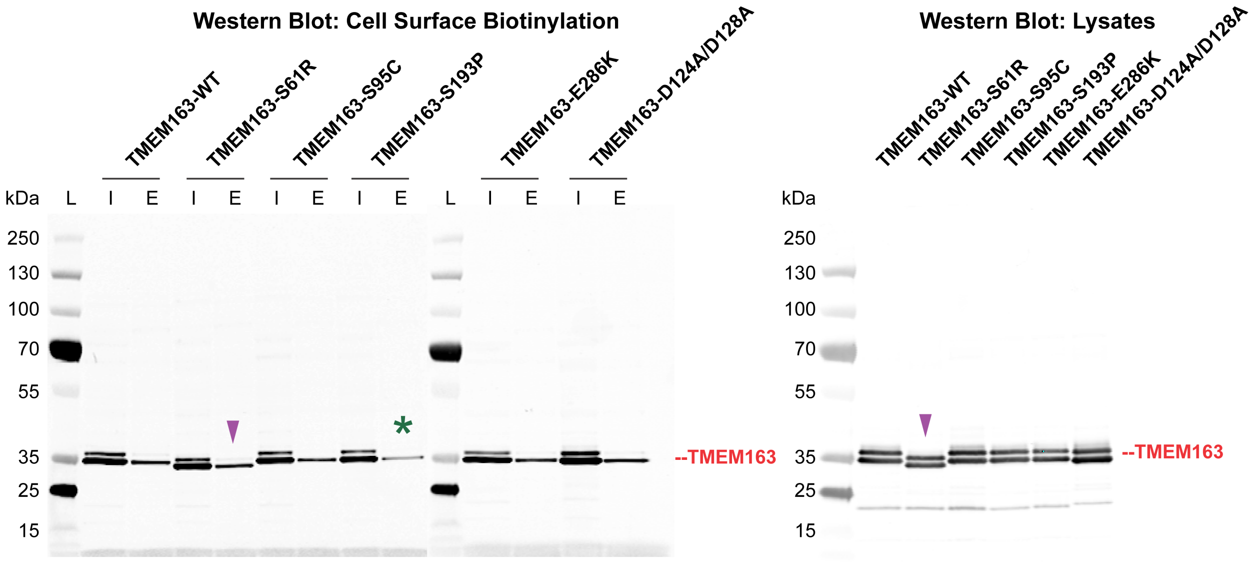


**Figure S5. Cell surface expression of wild-type TMEM163, TMEM163-D124A/D128A variant, and non-synonymous SNPs: S61R, S95C, S193P, and E286K.** Representative Western blot images of Myc-DDK peptide tag constructs probed with anti-DDK monoclonal antibody following CSB (*left panel*) and standard immunoblot of cell lysates taken from each expression construct (*right panel*). All TMEM163 expression constructs show protein localization within the PM. The PM localization of TMEM163-S193P (*green asterisk*) is noticeably reduced compared to other protein variants. This result complements the fluorescence microscopy data. The blots also show that TMEM163-S61R (*purple arrowhead*) migrated faster than the wild-type TMEM163 and other protein variants, possibly due to the arginine substitution of the serine residue. I, input; E, elution; (n = 3 independent trials).

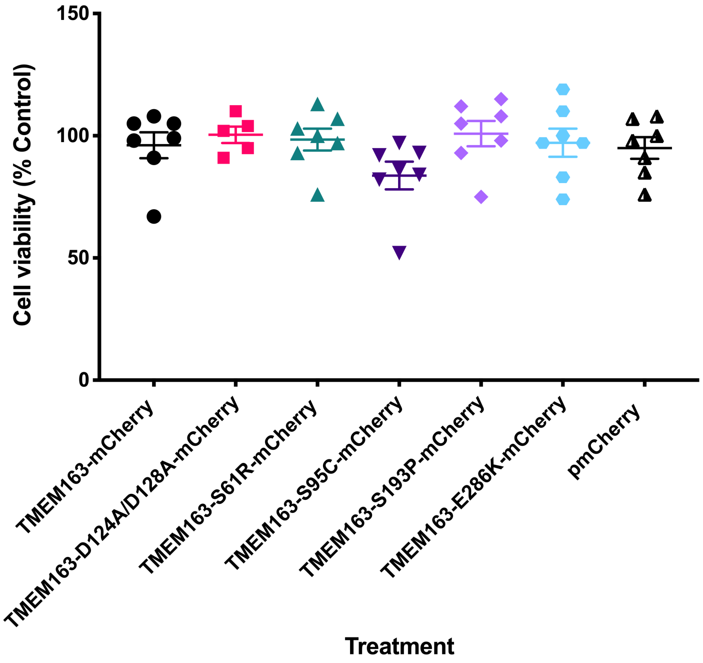


**Figure S6. Cell viability of HEK-293 cells expressing wild-type TMEM163, TMEM163-D124A/D128A variant, and non-synonymous SNPs: S61R, S95C, S193P, and E286K.** The mammalian expression construct contains the mCherry fluorescent protein tag. The TMEM163 protein variants do not significantly affect cell survival when compared to cells expressing the wild-type TMEM163 construct. Note, however, that cells expressing the S95C construct appear to have slight cytotoxicity in comparison with the other expression constructs. Data are represented as mean ± SEM of % control (n ≥ 4 independent trials).

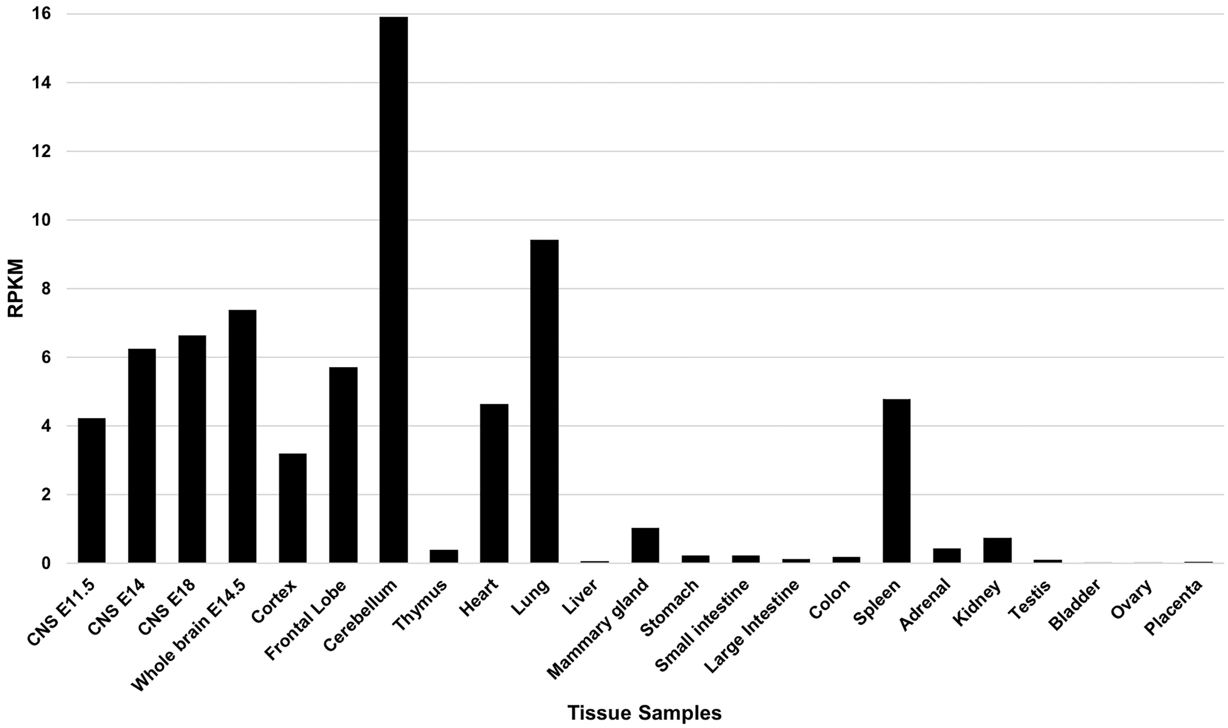


**Figure S7**. **Mouse Tmem163 tissue expression.** The expression of Tmem163 shows tissue specificity with relatively higher levels in embryonic (E11-E16) and adult central nervous system (CNS) tissues, especially in adult cerebellum and frontal cortex. Other tissues with relatively high levels include the lung, heart, and spleen, while relatively low levels are found in mammary and adrenal glands, thymus, and kidney. RPKM: reads per kilobase of transcript, per Million mapped reads. The RNA-seq data were obtained from the Mouse ENCODE transcriptome data project (Accession no. PRJNA66167) and the data were re-plotted using the Excel software.
